# Comparison of environmental, biological and anthropogenic causes of wildlife–vehicle collisions among three large herbivore species

**DOI:** 10.1101/385161

**Authors:** Saint-Andrieux Christine, Calenge Clément, Bonenfant Christophe

## Abstract

Wildlife–vehicle collisions are of increasing concern with regards to the continuous and accelerating anthropogenic development. Preventing and mitigating collisions with wildlife will require a better understanding of the environmental and biological drivers of collision risks. Because species of large mammals differ in terms of food requirements, habitat selection and movement behaviours we tested at the management unit level if the density of collisions with red deer, roe deer and wild boar differed in terms of spatial distribution and explanatory factors. From 20,275 documented collisions in France between years 1990 and 2006, we found marked differences in the most influential environmental factors accounting for the density of collisions among the three species. The effect of road density was higher for the red deer than for the two other species and did not level off at our spatial-scale of observation. As expected, the annual hunting harvest – interpreted as a proxy of population abundance – was positively associated with the density of collisions for all species, being the strongest for red deer. While the collision density decreased with the proportion of forest in a management unit for wild boar, it increased with the fragmentation of forests for red deer that commute among forest patches between day and night. To reduce the number of wildlife– vehicle collisions, our results suggest to generalise road fencing and/or a control of abundance of large herbivore populations. Mitigation measures should target units where the collision risk is the highest for the most problematic species.

## Introduction

Over the last decades, heavier traffic loads and continuous expansion of road networks paralleled the increasing number and distribution of most large herbivore populations (Milner et al. 2006, Massei et al. 2015), resulting in a dramatic increase of wildlife–vehicle collision frequency in many European and North American countries (Langbein et al. 2010). For large mammals between one to two millions wildlife–vehicle collisions are recorded annually in the United States (Huijser et al. 2015), and about a million in Europe (Langbein et al. 2010), generating acute ecological consequences on animal populations (Forman et al. 2003, Rytwinski and Fahrig 2015). The risk for human safety make wildlife–vehicle collisions an important socio-economic issue and a major issue in the road safety policies (Groot Bruinderink and Hazebroek 1996). The substantial economic bulk of wildlife–vehicle collisions and the need for mitigation of risks raised active areas of research (Schwabe et al. 2002, Malo et al. 2004, Huijser et al. 2015, Elmeros et al. 2011, Gunson et al. 2011, Hothorn et al. 2015). Understanding and identifying high collision risk areas and its ecological and biological drivers is of prime importance to predict wildlife–vehicle collisions in space and time, and to implement appropriate and efficient mitigation measures.

First and foremost, road infrastructure and use by drivers are anthropogenic explanatory factors potentially driving the occurrence of wildlife–vehicle collisions. The vehicle speed and the density of roads are since long recognized to increase the frequency of wildlife–vehicle collisions (Pojar et al. 1975, Case 1978, Hartwig 1993, Groot Bruinderink and Hazebroek 1996, Seiler 2005). For instance, the number of moose-vehicle collisions in Sweden was positively correlated with density of roads with a speed limit of 90 kph (Seiler 2005). Road class also influences the risk of collisions and mosts are recorded on secondary roads, mainly because they represent the greater cumulative length within most national road networks. Per unit of length, however, collisions frequency is higher on primary roads where traffic volume and vehicle speed are greater (Pojar et al. 1975, Bashore et al. 1985, Désiré 1992, Hartwig 1993, Hubbard et al. 2000, Langbein et al. 2010, Roedenbeck 2007, McShea et al. 2008). While the number of collisions increases broadly with traffic load, collision frequency can level off above a traffic density threshold, from which animals avoid crossing roads (Müller and Berthoud 1997, Skölving 1985, Clarke et al. 1998, Seiler 2004, 2005).

From an ecological point of view, space use and habitat selection by animals are key processes for the understanding of the spatio-temporal distribution of wildlife–vehicle collisions. Habitat selection by animals is a hierarchical process whereby choices observed at small scales are constrained by previous choices made at larger spatio-temporal scales (Johnson1980; Levin1992). Hence habitat characteristics, from landscape to the vicinity of the road, should influence wildlife–vehicle collisions at different spatial scales (de Bellefeuille and Poulin 2003). Most previous studies investigating collision patterns did so either at very large, state-wise or continental scales (Brockie et al. 2009, Červinka et al. 2015, Seiler et al. 2004), or at a very fine scale (Taylor and Goldingay 2004, Grilo et al. 2009). However, intermediate spatial scales are also relevant because in a patchy landscape collisions are more likely to happen on road sections located between woods and open fields because animals move frequently between protected resting areas and meadows or agricultural crops to forage (Puglisi et al. 1974, Bashore et al. 1985, Hubbard et al. 2000). This is the case for wild boars (*Sus scrofa*) commuting on a daily basis from forested patches to open fields (Carbaugh et al. 1975, Waring et al. 1991, Keuling et al. 2009). Consequently, when roads run through homogeneous landscapes, wildlife–vehicle collisions are more uniformly distributed in space than in fragmented and heterogeneous landscapes (Bellis and Graves 1971, Bashore et al. 1985, Hubbard et al. 2000). A study on white-tailed deer (*Odocoileus virginianus*) illustrates how landscape affect collision risks, being higher with the proportion of woodland cover level (Finder et al. 1999, Hubbard et al. 2000, Roedenbeck 2007). Matching observation with management scales of species is also way to provide managers with efficient policies they can act on, particularly because our ability to predict collision location increases with the spatial scale (*e.g*. Orrock et al. 2000).

Most large herbivores in Europe and North America are forest-dwelling species that are strongly attracted to roadsides (Bellis and Graves 1971, Carbaugh et al. 1975, Bashore et al. 1985, Groot Bruinderink and Hazebroek 1996). By regularly feeding on roadsides animals put themselves at a greater risk of collisions with vehicles (Puglisi et al. 1974, Bashore et al. 1985, Finder et al. 1999, Roedenbeck 2007), previous works searched for causal factors of wildlife–vehicle collisions, but focused on one single species (moose *Alces sp*.: Seiler (2005), Dussault et al. (2007); white-tailed deer: Bashore et al. (1985), Finder et al. (1999), Hubbard et al. (2000); roe deer: Mysterud (2004)), or did not differentiate among species (Malo et al. (2004), for red deer *Cervus elaphus*, roe deer *Capreolus capreolus* and wild boar, Gunson et al. (2009) and Nielsen et al. (2003), for white tailed deer and mule deer *Odocoileus hemionus*). In spite of many large hervivore species live sympatrically, comparative analyses of wildlife–vehicle collisions in mammals have rarely been conducted. Differences in diet and body size, space use behaviours, sensitivity to human presence and disturbance, or levels of grouping patterns (Sáenz-de Santa-María and Tellería 2015) could generate contrasting spatio-temporal distribution of collisions (as documented by Rodríguez-Morales et al. 2013). Our ability to predict where and when wildlife–vehicle collisions most likely occur may actually be hampered by specific habitat choice behaviours and its ecological correlates in the landscape.

Based on data recorded at the management unit (MU) scale over 9 departments between 1990 to 2006 in France, we first describe the spatial distribution of vehicle-wildlife collisions of the sympatric red deer, roe deer and wild boar. At this scale of observation, we aim at explicting what factors affect the number of roadkills in a given MU. To do so, we compare the relative effects of a set of environmental, biological and anthropogenic variables on the number of wildlife–vehicle collisions among species, using a Bayesian statistical framework to test the following predictions (Table 1):

**Table 1.** Hypothesis-predictions table. Here we present the rationale for the different predictions stemming from the hypotheses that the spatial distribution of wildlife-vehicle collisions differ according the species’ habitat selection behaviour. Symbols within brackets stand for the direction and relative magnitude of the effects for relatively to each of the three species of large herbivore consider in the study.

| Hypothesis | Red deer | Roe deer | Wild boar |
| --- | --- | --- | --- |
| Abundance | (++) | (++) | (++) |
| Habitat fragmentation | (0), present in forested areas mostly, excluded by hunters in other habitats | (+++), favourable environment | (++), Ubiquitous, wild boar use open areas to feed and forested areas for protection |
| Proportion of forested area in the management unit | (+++), use forested area mostly to minimize disturbance and is highly mobile | (++), roe deer favor fragmented habitats | (+), (Keuling et al. 2009) |
| Proportion of agricultural land in the management unit | (+), red deer uses agricultural crops close to forested areas | (+), roe deer favor fragmented habitats | (+++), (Keuling et al. 2009) |

1. We predict a positive relationship between the number of wildlife–vehicle collisions and the density of roads (Romin and Bissonette 1996, Pokorny 2006, Vignon and Barbarreau 2008), and between collision number and average car speed, hence leading to an increasing risk from local roads to highways. Also, the average number of wildlife–vehicle collisions should increase with population abundance, both across species and increasing from red deer, roe deer to wild boar, and in time with the annual variation of each species abundance (Schwabe et al. 2002, Seiler 2005, Sudharsan et al. 2006).
2. We expect more collisions with the proportion of forest in the landscape (Carbaugh et al. 1975, Waring et al. 1991) because red deer, roe deer and wild boar are all forest-dwelling species and found at highest densities in this habitat type (*e.g*. Telleria and Virgós 1997, Virgós 2002). However, the association between herbivore abundance and forest densities differs among the three species (Hewison et al. 2001, Patthey 2003, Saïd et al. 2005, Keuling et al. 2009, Thurfjell et al. 2009). In France red deer is confined by hunters to forests or to mountain areas, wild boar is also a forest species, but is currently expanding in all in all ecosystems thanks to its flexible behaviour, and roe deer is more ubiquitous and present in all departments (Maillard et al. 2010). Consequently, we predict a decreasing effect of the proportion of forest on the number of collisions from red deer, to roe deer and wild boar (Table 1).
3. Large herbivore move between different habitat patches so we expect landscape fragmentation to increase these movements and, as a consequence, the number of collisions too (Bashore et al. 1985, Romin and Bissonette 1996, Hubbard et al. 2000, Madsen et al. 2002). Moreover, with smaller home ranges, landscape fragmentation is less likely to increase an individual home range heterogeneity and among-patch movements. We hence expect the effect of habitat fragmentation on the number of collisions to decrease in magnitude from roe deer, wild boar and red deer following their respective average home range size (Table 1).
4. For large herbivores, crops are highly attractive food resources and individuals frequently commute between forests and agricultural areas on a daily basis (Keuling et al. 2009). We predict a positive association between the proportion of agricultural areas and the number of collisions but expect different responses for the three species with a stronger effect for wild boar and roe deer than for the red deer (Table 1). Wild-boar indeed use agricultural crop intensively (87% of the total amount paid for big game damage are done by wild boar, Maillard et al. 2010) and roe deer has colonised the agricultural plain (Hewison et al. 2009). Conversely red deer spatial distribution is restricted and strongly associated with forest (Milner et al. 2006), while being less attracted to crops than wild boar (Schley and Ropper 2003, Gebert and Verheyden-Tixier 2001).

## Material and methods

### Study sites

The local hunting associations of 9 French departments (Cher, Jura, Loire, Loiret, Moselle, Oise, Rhône, Haute-Savoie,Vendée; see Fig. 1 and Table S1 for a detailed description) collected and centralized the collision data. Despite our choice of the departments being primarily motivated by data availability, the 9 locations are representative of most mainland French ecosystems. We did not contribute to data collection ourselves and wildlife–vehicle collision cases were reported by the car driver or by direct observations of carcasses on the roads by officials (game wardens, police…). The monitoring spanned between 1990 and 2006 but varied among departments. Each department is divided into management units (MUs) defining administrative subdivisions of departments where game management is comprehensive and homogeneous. MU border may differ for red deer, roe deer and wild boar. Overall, we had 266 MUs in the 9 departments for roe deer, 247 MUs for wild boar in 8 departments (no wild boar data in Rhône), and 110 MUs for red deer in 7 departments (no red deer in Loire and Rhône). On average the surface of a single MU was 208 km2 (SD = 167 km^2^). Because the exact location of the collision was unknown (no GPS fixes), we assigned each collision event to the closest MU. Therefore potential location inaccuracies were of limited consequences on the presented results.

### Spatial scale of observation

Our statistical unit was the MU making the spatial scale of investigation of wildlife-vehicle collision pattern rather large. Because of this particular sampling design, we did not attempt to explain the location of collisions at a very fine scale *e.g*., by comparing local conditions where the collisions took place and a couple of meters away (case-control design, *e.g*. Eberhardt and Thomas 1991). Instead, we explain the collision number of each MU with the mean value of the different environmental variables measured across the corresponding MU to test their statistical association and to guaranty that number of MUs and environmental descriptors were of the same dimension. As for all processes of habitat use (Johnson 1980, Dupke et al. 2017), predictors of wildlife–vehicle collisions likely change with the spatial scales. At large spatial scale, habitat selection is related to landscape spatial structure, such as topography or habitat fragmentation. For example several authors have shown that vehicle collisions with red deer, roe deer and wild boar were more likely in forested environments at large spatial scale while at a smaller spatial scale, road sections with the highest collision risks were located in the open areas or at the forest border (Désiré 1992), or had roadsides with dense vegetation for roe deer (Madsen et al. 2002). In addition, the previously described barrier effect on road traffic and density on the number of collisions are not expected at large spatial scale of investigation because the range of road density is limited in each MU (see discussion).

### Explanatory variables of observation

Three types of explanatory variables were used to describe the road characteristics (anthropogenic variables), the landscape patchiness based on habitat composition (environmental variables), and the large herbivore populations (biological variables).

#### Anthropogenic variables

We described the road network using the Routes 500 database from the Institut Géographique National (IGN 2001) to derive the road density of MUs. We classified roads in four categories based on the importance of road sections for the traffic (see Supporting information 1): local roads, regional roads, national roads and motorways. For each road type, we assigned one of the three possible road density classes (low, medium and high) to the MU to explore non-linearity in the effect of traffic on wildlife–vehicle collisions. Because the statistical distribution of the road density is strongly asymmetrical, we had to find a statistical transformation to ensure the numerical stability of our results and avoid strong leverage effects. We set the limits of road density classes so that MU sample sizes were balanced in each class by computing the 1/3 and 2/3 quantiles of road density distributions. The limits defining the density classes differed according to road type. These limits were equal to 0.37 and 0.52 km per 100 ha for local roads (low, medium and high categories corresponded to road densities of <0.37, between 0.37 and 0.52, and > 0.52 km per 100 ha respectively), 0.16 and 0.24 km per 100 hectares for regional roads, 0.09 and 0.16 km per 100 hectares for national roads, and 0.01 and 0.03 km per 100 hectares for motor-roads. The Peason’s correlation (*ρ*) between pairs of variables measuring the density of roads never exceeded 0.36, so that we did not considered these variables as redundant. Note, however, that most roads in a MU are local roads, so that the overall road density is strongly correlated with the local road density (*ρ* = 0.72), and we did not try to test the effect of the overall road density alone, in addition to other road effects.

We also characterized roads sinuosity for each MU by calculating the ratio between the curvilinear length of the road segments and the distance in a straight line between the extreme points of the road. A straight road would have a sinuosity index of 1. We calculated the sinuosity only for local and regional roads because national roads and motorways were mostly straight. Fencing is an efficient way to reduce wildlife–vehicle collision risk (Clevenger et al. 2001, McCollister and Van Manen 2010), accounting for its confounding effects on collision density would be relevant. However, because the information about road fencing is not available nor centralized in France, we could not assess the effect of road fencing on collision density in our study.

#### Environmental variables

We extracted landscape variables with a GIS by mapping all MUs and calculating 10 descriptors of the environment. We used the CORINE Land cover (2006) database to derive a categorical variable “Habitat type” (HT) describing the land cover type in the MUs. For each MU, HT returns the proportion of the area covered by forest, agricultural crops, natural open areas, and urban and anthropogenic habitats (4 habitat types).

Moreover, we indexed the fragmentation of the forested habitat with the number of connected forest patches in the MU. Let *F*_*f*_ be the number of connected forest patches. To account for the larger number of forested patches in larger MUs we scaled *F*_*f*_ by the area of the corresponding MU, yielding a density of connected forest patches *D*_*f*_. However, because the relationship between *D*_*f*_ and the proportion of forest in the MU was non-linear (*i.e. D* was redundant with the land cover variable “Forest”), we used a nonparametric loess regression (degree = 2, span = 75%, Cleveland, 1993) to predict the logarithm of *D* as a function of the proportion of forest in a MU. Ultimately, we used the residuals of this regression as an index of the forest fragmentation (noted *R*_*f*_), whereby a positive values meant more forest patches in the MU for a given forest cover, and conversely for negative values. Following the same procedure, we calculated a fragmentation index of urban patches (*R*_*u*_), using the density of urban patches *F*_*u*_ instead of the density of forest patches. Finally, we used the IGN geographic database to calculate a 3-classes categorical variable of elevation defined as the proportion of the MU area found <600 m, between 600 m and 1 500 m, and > 1 500 m a.s.l.

#### Biological variables

For the three species of large herbivores, we assessed population abundance with the number of harvested animals per km^2^ for each MU (referred to “hunting bag”; see Seiler (2005), Morelle et al. (2013) for a similar approach). We used the number of harvested animals per km^2^ during the previous hunting season, spanning from September of year *t* − 1 to the end of February of year *t*, to characterise the abundance of a species during year *t* and to use it as a predictor of the density of wildlife–vehicle collisions. We hence make the assumption that annual hunting bag is positively associated with the population abundance of large herbivores. These data were provided by the local hunting associations in the 9 departments, for every hunting season for which we had collisions data.

### Bayesian model fit and variable selection

For each species, we first modelled the number of collisions in a MU with a log-linear model with mixed effects. We assumed a Poisson distribution for the number of collisions, and we classically modelled the logarithm of the mean of this distribution as a linear combination of the characteristics described in the previous section (the usual log-link was used to ensure a positive predicted mean number of collisions). We accounted for the overdispersion in the response variable by including Gaussian residuals in our linear predictor, following the approach of Hadfield (2010). We entered the department (9 levels factor) as a random effect on the intercept. We fitted this model in a Bayesian context with the JAGS software (Plummer 2016).

We used a Bayesian variable selection approach to identify the variables affecting the density of vehicle-wildlife collisions in a MU, that is the number of collisions per surface unit. More precisely, we implemented Kuo and Mallick (1998)’s method to estimate the probability that each variable influenced the mean number of collisions. The Kuo-Mallik’s approach also allowed to identify the best models predicting the number of collisions: we could estimate the probability of each possible model to be the best one describing our data (*i.e*. every possible combination of the variables describing the management unit), and select the most likely one. We checked MCMC convergence and good mixing of MCMC chains graphically. We also checked the convergence of the chain with the criterion of Raftery and Lewis (1992). This criterion indicated no lack of convergence for any parameter of any model (the minimum number of iterations required to allow the calculation of 95% credible intervals with an accuracy of 0.02 and a probability of 0.9 was much smaller than the 500 000 iterations of the chains for all models and all parameters). Overall, the goodness of fit of the models was excellent for all species. Technical details and a formal description of this approach are available as *Supporting information 2*.

Note that we could not use these models to compare the influence of a given predictive variable on the collisions across species. Indeed, the slope associated to a given variable in a regression model cannot be compared across models containing different variables and different sampling units (Becker and Wu 2007). We needed to fit a more general Bayesian model to allow for this comparison. We first focused on the MUs containing all three species. Then, we identified the set *D* of variables belonging to the best model identified by the Kuo and Mallick (1998)’s approach for at least one of the three species. Then, for a given species, we predicted the average number of collisions per unit area and per year not only as a function of the variables identified as important for this species, but also as a function of the variables identified as influential for the other species. We also fitted these models by MCMC, using the same approach as for the fit of the previous models (more formally, we replaced the set *B* by the set *D* in equation (1) of *Supporting information 2*), but focused on the interaction term coefficient to make inference on among species differences.

## Results

For all years and departments, we recorded 20,275 collisions for all species among which 69.9% were roe deer, 3.1% red deer and 27% wild-boar. The collision number averaged 6.37± 8.62 per 100km^2^ for roe deer, reaching up to 80 collisions per 100km^2^ for some MUs (see Fig. 2). For red deer and wild boar, the density of collisions was lower, with on average 0.93 ± 1.89 and 4.12 ± 8.09 per 100km^2^, following the same order as their respective relative densities.

### Red deer

The forest fragmentation and the hunting bag had the strongest effect on the density of collisions with the red deer (Table 2). Note that the model with the largest probability was characterized by slightly more than one chance out of two to be the true model (58%, see Table 3), indicating a significant uncertainty in the model selection process. Alternative models often included a measure of human density (*e.g*. density of local roads – second best model, density of national roads – fourth best model, fragmentation of urban areas – 5th best model). However, none of these alternative models was characterized by a large probability to be the best model, suggesting that the frequency of collisions between red deer and vehicles are essentially determined by the density of red deer, as well as the forest fragmentation as we expected from its behaviour. This best model indicated that the number of collisions was larger when the hunting bag was high and the forest was strongly fragmented (Fig. 3).

**Table 2.** Description of the variables and their associated probability of inclusion in the best model predicting the average density of vehicle-wildlife collisions (*n* = 20 275) for red deer, roe deer, wild boar recorded in 9 departments (administrative boundaries) of France between years 1990 and 2006. These probabilities (corresponding to *P* (*α*_*j*_ = 1), using the notation introduced in the text) were calculated using the Bayesian approach suggested by Kuo and Mallick (1998), introduced in *Supporting information 2*). We fitted statistical models separately for each species.

| Variable | Code | Red deer | Roe deer | Wild boar |
| --- | --- | --- | --- | --- |
| Elevation | elev | 0 | 0 | 0 |
| Urban areas fragmentation | urbFrag | 0.05 | 0.01 | 0.01 |
| Forest fragmentation | forFrag | 0.87 | 0.04 | 0.01 |
| Habitat type | hab | 0 | 0 | 1 |
| Hunting bag | hunt | 1 | 1 | 1 |
| Roads sinuosity | sinus | 0.03 | 0.01 | 0.01 |
| Density of local roads | locr | 0.13 | 0.14 | 0.01 |
| Density of regional roads | regr | 0.02 | 0.01 | 0.01 |
| Density of national roads | natr | 0.12 | 0.66 | 1 |
| Density of motorways | motr | 0.03 | 0.24 | 0.03 |

**Table 3.**
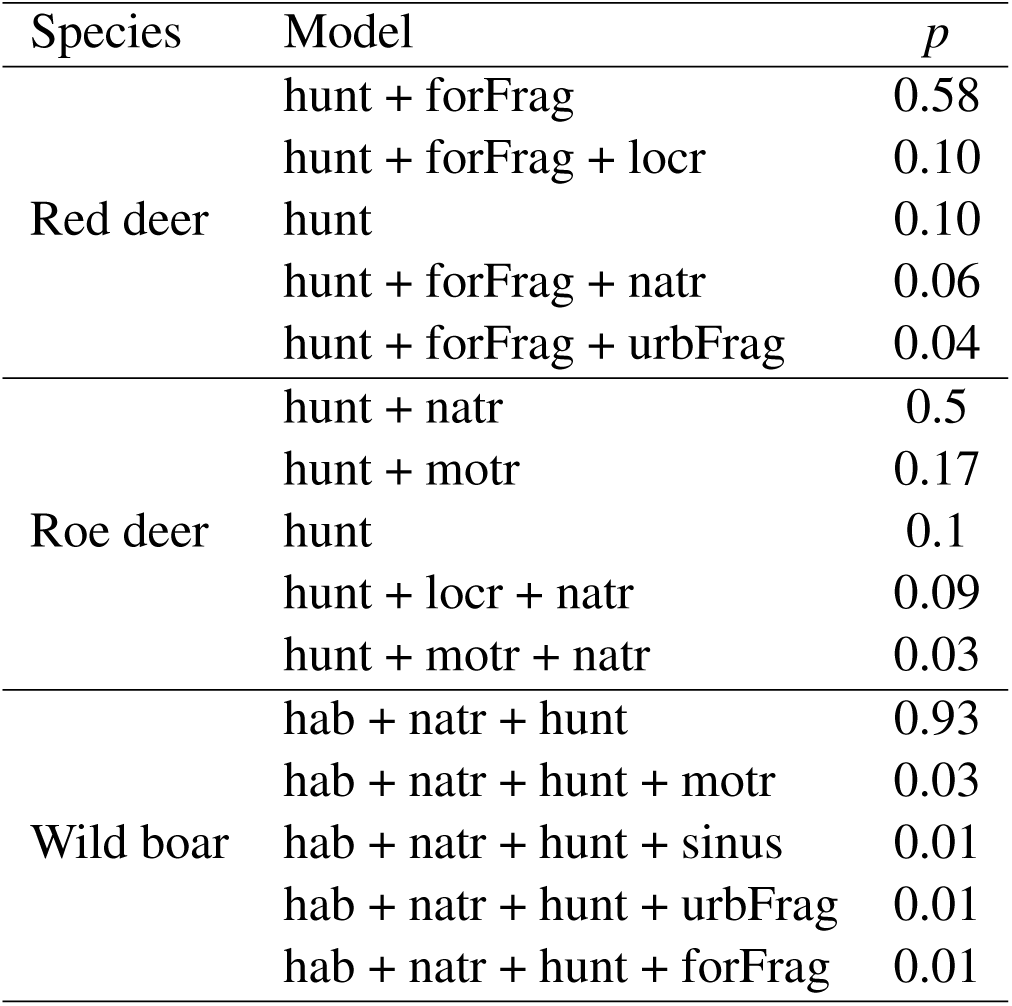
Probability (*p*) that the model is true, for the five best models predicting the average density of vehicle-wildlife collisions (*n* = 20 275) for red deer, roe deer, wild boar recorded in 9 departments (administrative boundaries) of France between years 1990 and 2006. These probabilities were calculated using the Bayesian approach suggested by Kuo and Mallick (1998), see *Supporting information 2*. The probability of a given model correspond to the proportion of Markov Chain Monte Carlo (MCMC) iterations for which *α*_*j*_ = 1 for the variables *j* of the model, and *α*_*j*_ = 0 for the other variables *j*′ (see *Supporting information 2* for a description of the parameters *α*_*j*_). See Table 2 for a description of the variable codes.

### Roe deer

Roe deer relative abundance and, to a lesser extent, the density of national roads were the main variables influencing the density of collisions with the roe deer (Table 2). Note that there was also a large uncertainty in the model selection for this species (Table 3). Although the best model was the model containing the two aforementioned variables, nearly all the other models with some statistical support included both roe deer relative abundance and one or several measures of road density (whether local roads, national roads or motorways). We fit the best model to describe the relationship between these two best variables and the density of collisions for the roe deer. This model indicated that the density of collisions was larger when both roe deer hunting bag and the density of national roads were high (Fig. 4). The effect of national road density was, however, small in comparison to the effect of the relative abundance of roe deer (Fig. 4).

### Wild Boar

Three variables strongly influenced the number of collisions between vehicles and wild boars: the habitat type, the hunting bag and density of national roads (Table 2). Note that there was only a small uncertainty on the model selection process as a very large probability is associated to this model (Table 3). We therefore fitted this best model (Fig. 5). To interpret the effect of habitat type, we had to account for the difference in average landcover between habitat types. For instance, while a MU with 80% of its area covered by agriculture is frequent in our data, a MU with 80% of urban area never occured in our data, so that we could not readily interpret the raw estimates of the coefficients to identify the differences between these habitat types clearly. To interpret these results, we predicted the average density of collisions between vehicles and wild boars as a function of the hunting bag and the density of national roads in four types of management units (see Fig. 5) in four contrasting environments: (*i*) a MU (labeled “Forest”), characterized by 91% of forest and 3% of the three other habitat types, (*ii*) a MU with a high land-cover of agricultural areas (labeled “Agri”), characterized by 91% of agricultural areas and 3% of the three other habitat types, (*iii*) a MU (labeled “Open”), characterized by 37% of open areas, 30% of forests, 30% of agricultural areas, and 3% of urban areas, and (*iv*) an urbanized MU (labelled “Urban”) characterized by 16% of urban area, 14% of open area, 35% of forests and 35% of agricultural areas. A careful examination of our dataset revealed that these particular sets of environmental conditions are typical of those encountered in France: thus, the habitat composition in the “Forest” MU is typical of the more forested management units observed in our dataset; the habitat composition in the “Urban” MU is typical of the highly urbanized management units, etc. According to the best model, the density of collisions increased with the hunting bag and the density of national roads (Fig. 5). On the other hand, the density of collisions was much lower in densely forested management units than in other types of management units.

### Comparison of patterns among species

We compared across species the effect of the variables identified by the Kuo and Mallik (1998)’s approach as important for at least one species. The coefficients of the variables for each species are given in Table 4. These models indicated that the effect of the forest fragmentation on the density of collisions was stronger for the red deer than for the two other species for which it was not different from zero. The effect of national roads was on average higher for the red deer than for the two other species, but this coefficient was characterized by a larger variance: the 90% credible interval included 0 for this species, explaining why this variable was not initally selected as part of the best model for red deer. The effect of the national road on the density of collisions was similar for the wild boar and the roe deer. The credible interval on the coefficients of the various habitat types (forest, urban, open, agriculture) included zero for all species except the wild boar. The density of collisions between vehicles-wild boar was negatively affected by the density of forest. Finally, relative abundance was positively related to the density of collisions for all species but was three times larger for the red deer than for the roe deer and wild boar.

**Table 4.** Coefficients of the environmental variables in the final model predicting the average density of vehicle-wildlife collisions (n = 20 275) for red deer, roe deer, wild boar. These collisions were recorded in 9 departments (administrative boundaries) of France between years 1990 and 2006. All fitted models include all the variables belonging to the best model for at least one species (identified by the Kuo and Mallik’s approach, see *Suporting information 2*). We present the 90% credible intervals in parentheses (intervals with a probability equal to 0.9).

| Variable | Red deer | Roe deer | Wild boar |
| --- | --- | --- | --- |
| Percentage of forest habitat | -2.44 (-9.64 – 4.76) | -1.1 (-7.21 – 4.53) | -6.17 (-12.31 – -0.33) |
| Percentage of agricultural lands | -6.81 (-14 – 0.31) | -0.69 (-6.87 – 4.98) | -4.48 (-10.71 – 1.57) |
| Percentage of open habitat | -4.32 (-11.54 – 2.85) | -1.43 (-7.69 – 4.34) | -3.95 (-10.22 – 2.17) |
| Percentage of urban habitat | -0.32 (-9.08 – 8.45) | -1.09 (-7.28 – 4.81) | 1.62 (-5.24 – 8.33) |
| Density of national roads | 0.38 (-0.02 – 0.78) | 0.21 (0.04 – 0.38) | 0.29 (0.04 – 0.53) |
| Hunting bag | 0.015 (0.009 – 0.022) | 0.005 (0.003 – 0.006) | 0.007 (0.006 – 0.009) |
| Forest fragmentation | 0.83 (0.52 – 1.18) | 0.1 (-0.03 – 0.23) | 0.03 (-0.15 – 0.2) |

## Discussion

By comparing the spatio-temporal pattern of vehicle collisions with red deer, roe deer and wild boar we found that population density and movement behaviour of animals govern collision density differently for the three species. Hunting bag, used as a proxy of population density, was the most influential variable, confirming that the species abundance in an area is positively associated to collision risks (Puglisi et al. 1974, Farrell et al. 1996, Seiler 2004) but this effect was, for instance, 3 times larger for the red deer than roe deer and wild boar. Similarly, for red deer and wild boar collisions with vehicles likely increase with movements between patches of favored habitats in response to habitat selection for foraging and disturbance. Conversely, collisions between roe deer and vehicles appeared more uniformly distributed in the landscape.

### Population abundance and collisions with vehicles

It has been repeatedly shown that local densities of large herbivores strongly influence collision risks (Puglisi et al. 1974, Farrell et al. 1996, Seiler 2004). Accordingly, we found that hunting bag, a proxy population abundance, is the only common predictor of wildlife–vehicle collisions for the three large herbivore species in agreement with our predictions. The effect is always positive but small as for a given MU, collision density increased by approx. 1% (range 0.5 – 1.5) for each additional thousands of animals shot per year. The relationship between population abundance and frequency of collisions differed among the three species, the largest effect being observed for red deer and the smallest for roe deer (Table 4). Accordingly, the annual number of wildlife–vehicle collisions increased 6-fold between 1986 and 2006 at the national level in France, while at the same time the hunting bag, a large-scale proxy of animal abundance, was 4 times larger for red deer, 5 times for roe deer and 6 times for wild boar (Vignon and Barbarreau 2008).

We contend, like previous studies (Iverson and Iverson 1999, Morelle et al. 2013), that we used hunting bag as a proxy of population abundance to account for wildlife vehicle collision. In France, annual quotas set how many roe deer and red deer may be harvested during the hunting season, while wild boar hunting is either unlimited or with quotas, which could also explain the different relationships between population abundance and collision density we find. Because red deer face high hunting pressure to control its colonisation to new areas and its associated damages on crops and forests, any increase in hunting bags likely reflects an increase in abundance (Saint-Andrieux et al. 2004). Conversely roe deer is ubiquitous in France, over a much longer period of time than red deer, and its population dynamics is currently levelling off (Maillard et al. 2010). Consequently, hunting bag may not capture variation in roe deer abundance as for the red deer, so that the relationship between hunting bag and collision frequency is weaker for the roe deer. Similarly, wild boar reproduction is strongly influenced by oak masting events (Gamelon et al. 2013), which leads to delayed effects on abundance not immediately reflected into hunting bags, and hence on its relationship with the number of collisions.

Clearly, hunting bag may be affected by hunting effort (Iijima 2017), the relative effect of relative abundance of herbivores on the number of collisions should be confirmed with more accurate estimates of population densities. Our results do not necessarily indicate a stronger effect of the animal population abundance on the density of collisions for the red deer. Alternatively, this difference could arise from a combination of different ecology, behavioural responses to hunting, and hunting practices among species.

### Importance of inter-specific differences

Related to their body size and specific food requirements, the ranging behaviour and abundance of the three species differ markedly. At the individual level, a large home range size increases the probability of road inclusion within and individual’s area of use. At the population level, species with the larger home range put more individuals at risk of collision than species with smaller home range. Having more individuals at risk of collision could in turn lead to a stronger association between population abundance and the density of wildlife–vehicle collisions. In mammals, home range size is associated with species’ body size (Lindstedt et al. 1986, Mysterud et al. 2001) and should decrease from red deer to wild boar and roe deer. Similarly in large herbivores population abundance is lower for large than small sized species (Silva and Downing 1995). Although limited to three species, the effect of hunting bag on collision frequency decreases from red deer, to wild boar and roe deer (Table 4), as expected from their respective average body size. The lower collision density we observe for red deer could result from a greater sensitivity to anthropogenic activities and avoidance behaviour of roads (Frid and Dill 2002). We currently have no direct observation from animal relocations to support this hypothesis and empirical evidence for road avoidance behaviour in large herbivores are equivocal. For instance, no road avoidance was found for red deer in Norway where traffic and road density are comparatively lower than in France (Meisingset et al. 2013). Conversely, D’Amico et al. (2016) reported road avoidance for red deer and wild boar although to a similar extent for the two species (see also Rowland et al. (2000) on elk in North America).

Our results show that although the risk of collision exists in all kinds of environment, it differed in magnitude according to the species and the landscape structure. Collisions with wild boars were more likely to occur in MU with agricultural, urban and open areas and less in forested areas, which differentiates this species from red deer and roe deer (Table 3; Fig. 3 & 4). The concomitant growth of urban areas and wild boar populations in recent years has increased the presence of this highly plastic species around cities and in suburbans areas (Cahill et al. 2012) where hunting is difficult and rather limited. As urban encroachment expands on the agricultural and forested lands, we expected more collisions with the wild boar in urbanized areas. Wild boar frequently move from one habitat to another and does not preferentially seek forest patches, explaining why habitat type rather than forest fragmentation accounts for collisions number for this species. Opposed to wild boar is the typical forest dwelling red deer which occasionally feeds on of agricultural and mixed habitats in the vicinity of forest patches. Males make large movements across seasons to commute between foraging and rutting areas (Hamann et al. 1997), which put them at risk of collision with vehicles. In lowland forests, females have limited seasonal migration movements, but can travel relatively long distances to reach foraging areas at night (Klein and Hamann 1999). In agreement with the hypothesis that animal movement put them at risk of encounter with vehicles, we found that collisions with red deer were more numerous when the number of forest patches increases in the landscape (Table 4; Fig. 2). Conversely, the role of landscape structure had no detectable influence on roe deer collisions contrary to our predictions. Since roe deer is found in all habitats, population abundance could reflect habitat selection at the population scale better than our environmental variables. Moreover, habitat selection of ecotones by roe deer likely occurs at a small spatial scale and because they live on a small home range, forest fragmentation is less likely to increase home range heterogeneity and inter-patch movements. Consequently our environmental variables, measured at the management unit level, may not capture spatial heterogeneity relevantly for roe deer.

### Influence of road network on collisions

The road network configuration in the landscape is a key structuring factor of the collision risks with wildlife. A review of the wildlife–vehicle collisions with mammals showed that 39% of studies (7 out of 18) reported a positive effect of traffic, road width or speed limit on collisions number (Gunson et al. 2011). The “national” roads are the only type of roads affecting collision numbers with wild boars and roe deer in France (Fig. 3 & 4). Similarly in Slovenia, among a set of 40 tested variables describing landscape features, density of roads has an overwhelming effect on the number of collisions with roe deer (Pokorny 2006). These “national” roads connecting the main urban centres in France combines a heavy traffic load with a low level of protection such as fences or wildlife crossing and make most of the transportation network. Unexpectedly though, the density of roads has no detectable effects on collision number with red deer that was more related to landscape structure. Red deer behaviour in habitat selection could account for this relatively low the risk of collision. For instance, red deer select mostly for forested habitats with only a marginal use of agricultural crops (Hamann et al. 1997), hence restricting the area at risk of collision. In addition red deer forage on food patches close to roads at a time of low traffic burden mostly (Meisingset et al. 2013) or select habitat away from roads with the greatest vehicle use such as primary roads (Montgomery et al. 2012).

The density of other road types seems to have little influence on the density of collisions, whatever the species (Table 2). In France, highways and motorways are fenced most of the time. According to our results, road fencing proves to be an efficient method to limit collision risks despite a high speed limit (>110 km.h-1). Accordingly, in France in 2008 and 2009, 86 777 vehicles collisions with red deer, roe deer and wild boar have been recorded by insurance companies, of which only 1% occurred on highways and in spite of highways represent 97% of the daily traffic (40 400 versus 1 030 vehicles/day/km for highways vs other roads in 2010 respectively). Conversely, lower speed and narrower lane width likely limit traffic load on this secondary road network both factors being known to reduce wildlife–vehicle collision risks substantially (Hubbard et al. 2000).

We found more collisions with roe deer and wild boar in MUs with a higher road density and with a traffic ranging between 2 500 and 10 000 vehicles per day. More surprisingly collisions frequency kept on increasing with traffic over than 10 000 vehicles per day and we could not detect any barrier effect. Previous studies indeed suggested that traffic volume could prevent animals to cross the road Skölving (1985), Clarke et al. (1998), with varying thresholds (approx. 4 000–5 000 vehicle per day in Sweden: Seiler (2005); >10 000 vehicles per day in Germany: Müller and Berthoud (1997)), which is consistent with behavioural observation of habitat selection of red deer patterns according to traffic volume in red deer for instance (Meisingset et al. 2013). In our study, the traffic load of roads is likely confounded with road type and road fences, which may have hampered our ability to test for the effect of traffic volume *per se*, or to detect any threshold effect of traffic load on collision risk. Our large working spatial scale is an alternative explanation for the absence of saturation effect of traffic load on collision number. Being averaged over *>* 100km^2^, the range of road density values across MUs is limited. It is unlikely that the road density would be so large over a whole MU that the number of collisions would plateau in this MU.

### Management implications and conclusion

In spite of its economic burden (Bissonette et al. 2008) and ecological consequences (Forman et al. 2003), wildlife–vehicle collisions with large mammals are most often considered as a general, non-specific problem. Improving the efficiency of mitigation measures likely requires a better and finer knowledge about the causes of collision risks, including inter-specific variations. For instance in Spain, collisions with vehicles did not occur at the same time of the year or of the day, and at different geographical locations for roe deer and wild boar (Rodríguez-Morales et al. 2013). Our results suggest two lines of actions to mitigate wildlife–vehicle collisions. The species-specific factors affecting the collision process can help to focus the measures on high-risk areas, depending on which species is the most at risk in a management unit. For example, we have demonstrated that a high forest fragmentation can increase the collision risk with the red deer in management units where this species is dense. Thus, focusing the mitigation measures on roads crossing highly fragmented forests could reduce this risk. If most collisions occur with the wild boar in a department, the mitigation measures will be more efficient if focused on roads crossing areas where both the wild boar density is large and the agricultural habitat is large.

An alternative measure would be road fencing as most – but not all – highways and motorways, that are fenced in France, had lower collision density despite their high traffic load (between 1 and 2% of ungulates were killed on motorways: Clevenger et al. 2001, McCollister and Van Manen 2010). The number collisions with the red deer increasing with forest fragmentation, road fencing should be focused on roads crossing the most fragmented forests and targetting MU with high number of collisions with that species. Alternatively, fragmentation could be reduced with green bridges, overpasses and underpasses, in addition to maintaining connectivity for large herbivores (reduced connectivity is a major ecological consequence of road fencing, see Forman and Alexander 1998). Because vehicle speed increases collision density for wild boar and roe deer, speed should be reduced in high collision zones with these species. Alternatively, reducing population densities of large herbivores could limit the number of collisions per MU. Previous studies showed a reduction in the frequency of deer-vehicle collisions with lower deer densities (Rondeau and Conrad 2003, Sudharsan et al. 2006). Nevertheless, for population size reduction to lead to a substantial reduction in the collision frequency with wildlife would require a massive hunting effort, given the rather weak relationship we report here for the three species.

Awareness and prevention campaign could also be a way to mitigate wildlife–vehicle collision. Warning signs are often used in France to reduce wildlife vehicle collisions by warning drivers about the potential presence of wildlife on the road, although only the one sign pictures a jumping deer. The efficiency of the warning signs would be improved if adapted to the local risks with the appropriate species. For exemple in departments with no red deer and many wild boar, a warning sign with a wild boar would definitely be more relevant. An alternative way of reducing the number of collisions may be a better information of the motorists for whom collision risk mainly occurs on roads driving through forests. Information campaigns are needed for a general awareness that the collisions can take place everywhere, even around cities and on highways. Finally, motorists would benefit from a better knowledge about large herbivore behaviour. For example, if one wild boar crosses the road, a second animal or a third one is likely to come out.

## Acknowledgments

We thank the French Fédérations départementales des chasseurs of Cher, Jura, Loire, Loiret, Moselle, Oise, Rhône, Haute-Savoie et Vendée for providing us with the collisions data. We are grateful to two anonymous referees who greatly helped at improving previous drafts of the manuscript.

## Data and code accessibility

We have bundled the code used for the analysis as well as the complete data set in a R package named “ungulateCollisions”. This package contains a vignette named “modelfit” which describes how the analyses carried out in this paper were performed. To install it, the reader should first install the software JAGS on their computer (see http://mcmc-jags.sourceforge.net/for_instructions). The reader can easily install the package and access the vignette by copying the following R code in the R console:

~~~
library(devtools)
install_github(“ClementCalenge/ungulateCollisions”)
vignette(“modelfit”)
~~~

## Figure captions

**Fig. 1** Location of the nine departments (administrative subdivision of the country) from France used in our study, where *n* = 20 275 wildlife–vehicle collisions were recorded between 1990 and 2006.

**Fig. 2** Average spatial distributions of vehicle-wildlife collisions with with red deer, roe deer and wild board in each of the management units of the nine departments where collision data have been collected in France.

**Fig. 3** Model predictions of the average density of collisions between red deer and vehicles per year and per squared kilometre in a management unit, as a function of the hunting bag (x-axis) interpreted as a proxy of population abundance, and the forest fragmentation (from top to bottom). Wildlife–vehicles collision data were collected in 9 departments (administrative boundaries) of France from 1990 to 2006. To enable a graphical display of the fit, we considered three values of forest fragmentation: (a) low fragmentation (value = -1; only 2% of the management units present a value lower than -1), (b) medium fragmentation (value = 0, corresponding to the mean fragmentation observed in a management unit), and (c) high fragmentation (value = 1; 4% of the management units present a value larger than 1). The central curve corresponds to the point prediction of the model. The four shades of grey indicate (from darker to lighter shades): 20%, 40%, 60% and 80% credible intervals.

**Fig. 4** Model predicting the average density of collisions between the roe deer and vehicles per year and per squared kilometer in a management unit, as a function of the hunting bag (x-axis) interpreted as a proxy of population abundance, and the density of national roads (from top to bottom). Wildlife–vehicles collision data were collected in 9 departments (administrative boundaries) of France from 1990 to 2006. This latter variable was already discrete, so that we did not have to discretize it, as for figure 1. Relative roe deer abundance increases from panel a to c. The central curve corresponds to the point prediction of the model. The four shades of grey indicate (frow darker to lighter shades): The four shades of grey indicate (from darker to lighter shades): 20%, 40%, 60% and 80% credible intervals.

**Fig. 5** Model predicting the average density of collisions between the wild boar and vehicles per year and per squared kilometre in a management unit, as a function of the hunting bag (x-axis) interpreted as a proxy of population abundance, the density of national roads (increasing from top to bottom, this variable was already discrete as for figure 1), and the landcover by four habitat types. Wildlife–vehicles collision data were collected in 9 departments (administrative boundaries) of France from 1990 to 2006. We have defined 4 types of management units here (see text for details): (a, e, i) a forested management unit (“Forest”), (b, f, j) an agricultural management unit (“Agri”), (c, g, k) a management unit with an important landcover by open natural areas (“Open”), and (d, h, l) an urbanized management unit. The central curve corresponds to the point prediction of the model. The four shades of grey indicate (from darker to lighter shades): 20%, 40%, 60% and 80% credible intervals.

## Supporting information 1: Road classification and detailed description of the 9 departments of France

Road classification in France is official and based on its function, reflecting the size and administrative importance of the different urban areas it connects. The different functional types of roads has been defined by a document called “l’ARP, aménagement des routes principales”, available for download at http://dtrf.setra.fr/notice.html?id=Dtrf-0001919, published in 1994 and edited by the French Ministry of the Infrastructure. An updated overview of the different road types may be found in Wikipedia at https://fr.wikipedia.org/wiki/Classification_fonctionnelle_des_routes_nationales_en_France. Three main road types are currently recognized:

- *L* type road, long distance roads connecting major urban centers, mainly highways with two causeways;
- *T* type road, or transit roads, encompasses express roads with one causeway and two to three lanes;
- *R* type road, are multi-function roads, making most of the French road network. Those roads have various configurations ranging from two causeways for inter-city roads to one lane roads in the countryside;

The above mentioned road classification is the basis of the road network mapping available from the Institut Géographique National (IGN), the French Geographic Institution, in its ROUTE 500 database (http://professionnels.ign.fr/route500). We used the road classification by IGN in our paper, which subdivises the R road type into two more homogeneous road types. The road categorization by IGN strongly correlates with the average traffic density and road width. Our classification is ecologically relevant because roads more and more difficult to cross for wildlife from local to highway roads.

**Table S1.** Description of the 9 departments of France where the collisions between vehicles and wildlife have been recorded from 1990 to 2006. Column headings mean population density for ‘pop’, average elevation a.s.l. for ‘elev’, and minimum and maximum elevation a.s.l. for ‘elev. min’ and ‘elev. max’, the percentage area of forest for ‘% for.’, the road network density for ‘rd. dens.’, the percentage of motorways for ‘% Mways’, the percentage of national roads for ‘% nat.’, the percentage of regional roads for ‘% reg.’ and the percentage of local roads for ‘% loc.’. The last column describes the main land use and habitat types of each department.

| Department | area<br>(km<sup>2</sup>) | pop.<br>ind/km<sup>2</sup> | elev.<br>(m) | elev.<br>min. (m) | % for.<br>max. (m) | rd<br>dens. | % Mways | % nat. | % reg. | % loc | Land use and main habitats |
| --- | --- | --- | --- | --- | --- | --- | --- | --- | --- | --- | --- |
| Cher | 7 235 | 43 | 87 | 502 | 24 | 1.4 | 1 | 1 | 47 | 51 | Extensive crops, vineyard, deciduous forest, multiple cropping and grove |
| Jura | 4 999 | 52 | 177 | 1 495 | 44 | 2.1 | 1 | 1 | 33 | 65 | Lowland forest <200-300m a.s.l.; grassland plateau from 400 to 700 m a.s.l.; mountains and coniferous forest >1000 m a.s.l. |
| Loire | 4 781 | 159 | 130 | 1 631 | 26 | 2.6 | 1 | 1 | 30 | 68 | Continuous mixed forests of beech and fir tree, vineyards, moorlands |
| Loiret | 6 775 | 99 | 67 | 281 | 26 | 1.7 | 2 | 0 | 32 | 66 | Large agricultural lands, deciduous forest and typical peat swamp forest from Sologne |
| Moselle | 6 200 | 168 | 145 | 986 | 28 | 1.8 | 2 | 1 | 39 | 58 | 50% area is used for agricultural crops, mixed forest surrounds the Vosges mountain foothills |
| Oise | 5 860 | 140 | 22 | 236 | 21 | 2.1 | 1 | 1 | 33 | 65 | Vast cultivated lowlands, deciduous forests |
| Rhône | 2 715 | 166 | 140 | 1 008 | 22 | 2.1 | 2 | 1 | 27 | 70 | Vineyards, small crops, mid-elevation mountains covered with mixed forests |
| Haute-savoie | 4 388 | 181 | 250 | 4 809 | 39 | 2.1 | 2 | 0 | 33 | 65 | Grasslands, beech forest at low elevation and Alpine grasslands |
| Vendée | 6720 | 99 | 0 | 290 | 5 | 2.4 | 1 | 0 | 29 | 70 | Grove and coastal pine forests |

## Supporting information 2: Bayesian modelling fit and variable selection

We used a Bayesian variable selection approach to identify the variables affecting the most the density of wildlife–vehicle collisions in a MU. As a first step, we replicated the same approach for each species. For a given species, let *N*_*i*_ be the number of collisions with a vehicle in the MU *i*. We assumed that this variable could be described by the following over-dispersed Poisson distribution:

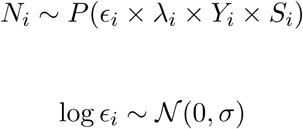

where *Y*_*i*_ is the number of years of data available in the MU *i, S*_*i*_ is the area of the MU *i, λ*_*i*_ is the average number of collisions per unit area and per year expected under our model (see below) and *ϵ*_*i*_ is a normal over-dispersion residual with zero mean and a standard deviation equals to *σ*.

The average number of collisions per unit area and per year in a MU *i* was modeled as a function of the *P* = 10 variables described in the last section (Table S1), according to the following log-linear model:

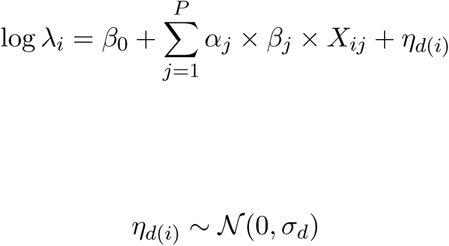

where *X*_*ij*_ is the value of the *j*th variable describing the MU *i, η*_*d*_ is a random effect describing the effect of the department *d, d*(*i*) is the department corresponding to the MU *i*, and *α*_*j*_ and *β*_*j*_ are two coefficients characterizing the role of the *j*th variable in this linear combination: (*i*) the coefficient *α*_*j*_ can only take values 0 and 1. When this coefficient is equal to 1, the *j*th variable belongs to the model; when this coefficient is equal to 0, the *j*th variable does not belong to the model. In a Bayesian context, the value of this coefficient is therefore considered as the realization of a Bernoulli variable characterized by a probability *p*_*j*_, which is the probability that this variable belongs to the model; (*ii*) the coefficients *β*_*j*_ can take any real value, and determines the importance of the *j*th variable on the average number of collisions when this variable belongs to the model, as in a classical regression model. This approach consists in separating the presence of a variable in a model from its importance, and then to estimate the probability of presence of each variable in the model from the data, as suggested by Kuo and Mallick (1998). In the rest of this paper, we refer to this approach as the Kuo and Mallick (1998)’s approach.

We set the following vague priors on the coefficients of the model:

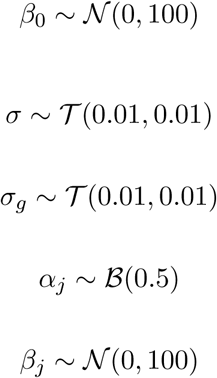

Posterior distributions of parameters were deduced from prior information about parameters and likelihood functions of the data by Monte Carlo Markov Chain (MCMC) simulations, i.e, inferences are made empirically by collecting many realizations from the posterior distribution using a variant of Metropolis method called Gibbs sampling (Gilks and Richardson 1996). We ran one chain for an initial period of 1,000 cycles (burn-in period) and then collected information for the next 500,000 iterations. We implemented the MCMC simulations with the JAGS software (Plummer 2010). From our analyses, we could (*i*) identify those variables with the largest influence on the number of wildlife–vehicle collisions and calculate the probability P (*α*_*j*_ = 1) that each variable *j* belong to the best model; (*ii*) identify the best models predicting the number of collisions and calculate the probability P(*α*_1_, *α*_2_, …, *α*_*J*_), for each possible combination of the coefficients {*α*_1_, *α*_2_, …, *α*_*J*_}, that the corresponding model is the best model. Then, for each species, we fitted and interpreted the best regression model predicting the average number of collisions per unit area and per year, *i.e*.:

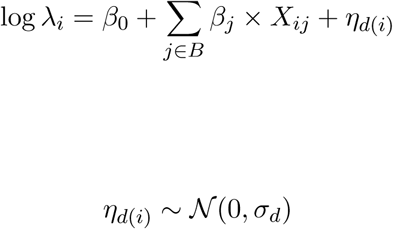

where *B* is the set of variables corresponding to the best model identified by the Kuo & Mallik’s approach. We checked the convergence of the MCMC chains both visually and by using the diagnostic of Raftery and Lewis (1992). None of these diagnostics showed any evidence of nonconvergence of the MCMC. We then examined the fit of the model using the approach recommended by Gelman and Meng (1996). For every iteration of the MCMC, i.e. for every value *θr* = *β*_0_ … of the vector of parameters of this second model sampled by MCMC, we simulated a hypothetical replication of the dataset using equation (1), i.e. we simulated a number of collisions in each MU. We then compared the observed number of collisions with the statistical distribution of simulated numbers of collisions. We calculated that 99% (red deer), 100% (roe deer) and 100% (wild boar) of the 90% of the credible intervals contained the observed number of collisions, which indicates that the fit was correct for the three species.

Note that it is difficult to compare coefficients *β*_*j*_ of a variable *j* across models containing different variables and different sampling units (Becker and Wu 2007), which precluded the comparison of the models across species. We needed to fit another general Bayesian model to allow for this comparison. We first focused on the MUs containing all three species. Then, we identified the set *D* of variables belonging to the best model identified by the Kuo and Mallick (1998)’s approach for at least one of the three species. Then, for a given species, we predicted the average number of collisions per unit area and per year not only as a function of the variables identified as important for this species, but also as a function of the variables identified as influential for the other species (more formally, we replaced the set *B* by the set *D* in equation (1)). We also fitted these models by MCMC, using the same approach as for the fit of the previous models. Note the collision data can be downloaded and the analyses replicated or compared with other approaches (see section Data and code accessibility).

